# Role of Caveolin 1 in metabolic programming of fetal brain

**DOI:** 10.1101/2022.10.18.512714

**Authors:** Maliha Islam, Susanta K Behura

## Abstract

Caveolin-1 (*Cav1*) encodes a major protein of the lipid rafts, called caveolae, which are plasma membrane invaginations found in most cells of mammals. *Cav1*-null mice, at an early adult age, exhibit symptoms that are hallmarks of Alzheimer’s disease, and show brain aging similar to that of one and half year old wildtype mice. In the present study, integrative analysis of metabolomics, transcriptomics, epigenetics and single cell data was performed to test the hypothesis that metabolic deregulation of fetal brain due to lack of *Cav1* influenced brain aging in these mice. The results of this study show that lack of *Cav1* deregulated lipid and amino acid metabolism in the fetal brain. Genes associated with the deregulated metabolites were significantly altered in specific glial cells of the fetal brain, and epigenetically altered in a coordinated manner with specific genes of mouse epigenetic clock. The interaction between metabolic and epigenetic changes in the fetal brain altered gene expression of the brain at old age. Together, these results suggested that metabolic deregulation in the fetal life elicited an epigenetic memory that altered brain programming for aging in *Cav1*-null mice.

## Introduction

Caveolin-1 is a major protein of the lipid rafts called caveolae which are omega shaped plasma membrane invaginations of most cells in mammals and other vertebrates. Mice lacking the Caveolin-1 gene (*Cav1*) are viable and fertile but show impaired endothelial functions (1), angiogenesis (2), and hyperproliferative and vascular abnormalities (3). These mice show reduced lifespan (4) and exhibit neuronal aging at young age (3-6 month old) that resembles brain of one and half years old wildtype (WT) mice (5). Multiple hallmarks of Alzheimer’s disease (AD) such as increased amyloid beta, tau, astrogliosis and shrinkage of cerebrovascular volume are observed in these mice at an early adult age (6). *Cav1* modulates beta-secretase required for the production of amyloid beta peptides in AD (7). Thus, loss of *Cav1* has been suggested to trigger AD pathologies in these mice upon aging (6).

Experiments with animal models of AD have shown that amyloid beta levels and gradual decline in cognitive ability are associated with abnormal molecular conditions of brain during early life (8–11). The ‘*Latent Early-life Associated Regulation’* (LEARn) model proposed by Lahiri and Maloney (12) suggests that changes in the epigenetic state of the brain during early-life serve as precursors for development AD pathologies later in life. While aging is the most influential factor of AD (13, 14), accumulating evidence suggests that specific stress or deficiencies during fetal stages are linked to increased risk of neurological disorders including AD (8, 15–20). Our recent study showed that aging of the brain is linked to early development of the brain via epigenetic influence of the placenta (21). Epigenetic memory is an emerging concept in fetal origin of adult heath and disease (22). Accumulating evidence suggest that epigenetic memory plays a key role to control effects of early-life environmental stresses on aging, and also to pass on parental exposures and experiences to the offspring (23–28).

Metabolic and epigenetic changes in the brain are associated with the risk of AD (25, 29–33). Abnormal level of urea, cholesterol, serine and other metabolites in the brain cholesterol increases risk of AD (34–37). Besides roles in the regulating urea (38) and serine (39), *Cav1* plays a role to regulate cellular homeostasis of cholesterol (7, 40–42). Loss of *Cav1* has been shown to deregulate cholesterol metabolism in embryonic fibroblasts and peritoneal macrophages in mice (43). Links between cholesterol homeostasis and epigenetic modification (44) have been shown in age-related diseases including cancer and dementia (33, 45–50). In this study, we show that lack of *Cav1* deregulates multiple lipids and amino acids, and genes associated with the metabolic pathways of these lipids and amino acids are impacted in specific glial cells of the fetal brain. Furthermore, we show that specific metabolism genes are differentially methylated along with epigenetic clock genes in distinct patterns in the fetal brain. These genes are significantly altered in the brain upon aging. Together, the findings of this study suggest that *Cav1* plays a role in the metabolic programming of fetal brain for aging.

## Results

### Metabolic deregulation of fetal brain due to Cav1 ablation

Untargeted metabolomics analysis showed that ablation of *Cav1* significantly (*p* < 0.02) deregulated several lipids and amino acids in the fetal brain (**Figure 1**). Cholesterol, hexadecanoic acid (known as palmitic acid), and steric acid, which are linked to Alzheimer’s disease (51, 52), decreased in the fetal brain of *Cav1*-null mice. While cholesterol and palmitic acid decreased by 10 and 4.7 folds respectively. Stearic acid decreased by 100 folds (**Table S1**). L-alanine and beta-alanine, which is produced by catabolism of hydroxyuracil, are essential for brain function (53–55) and are metabolically linked to cholesterol (54, 56, 57). We observed that L-alanine and hydroxyuracil decreased in the fetal brain due to the ablation of *Cav1*. However, several amino acids and organic compounds increased in the fetal brain in response to the ablation of *Cav1*. The alpha glycerophosphate ester, an esterified form of glycerol, increased by ~60 fold. Urea level, known to increase risk of Alzheimer’s and other forms of dementia (34, 58, 59), increased drastically by 2,183-fold in the fetal brain of *Cav1*-null compared to WT mice suggested that *Cav1* had a role in regulating brain urea which is independent of circulating urea (60).

**Figure 1.**
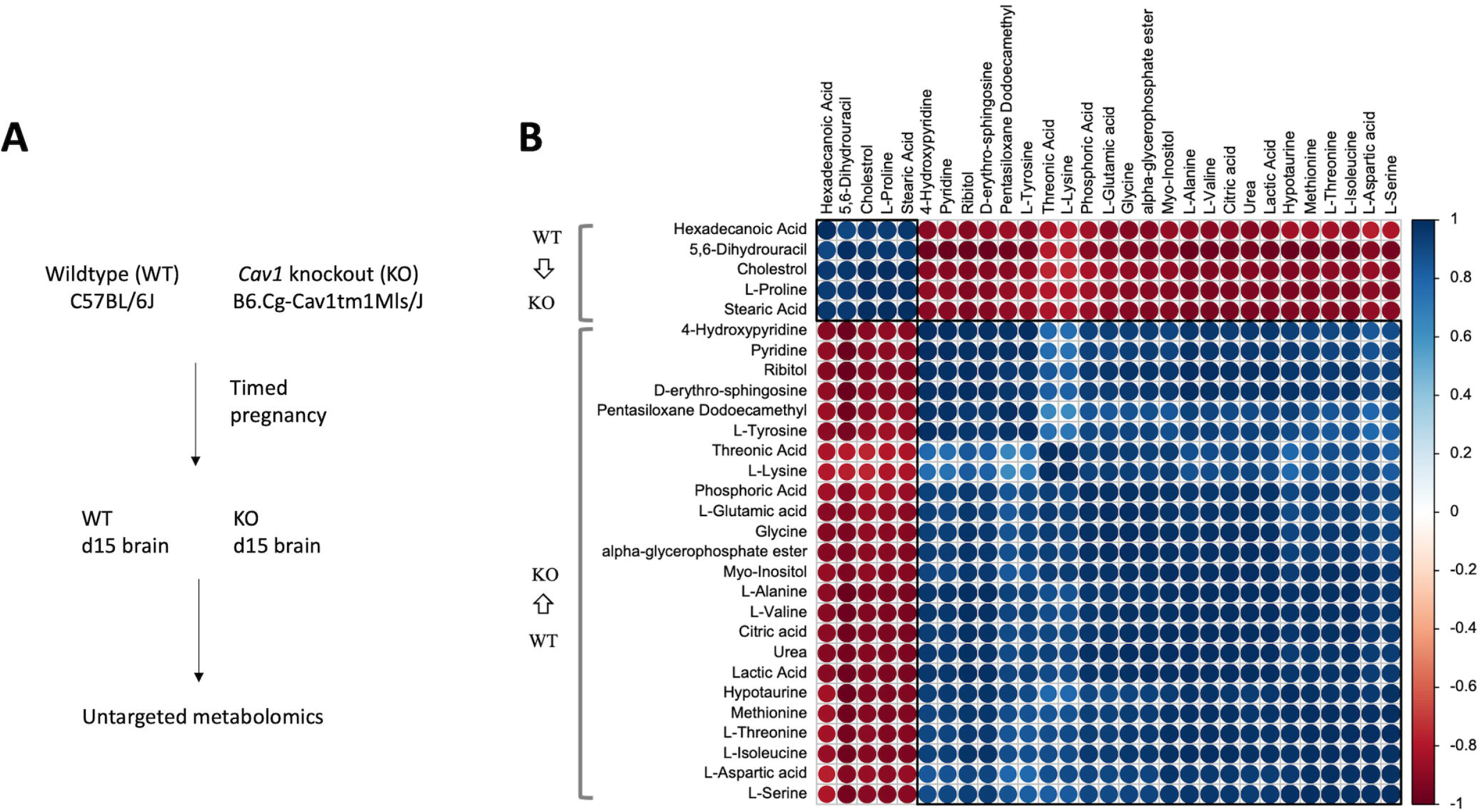
Experimental design and metabolomics. **A**. Timed pregnancy was performed separately with WT and *Cav1*-null mice to collect day 15 fetal brain for metabolomics (and other omics) analysis. **B**. Down-regulation and up-regulation of specific metabolites in the fetal brain. Pair-wise cluster analysis of variation of metabolite levels show that specific lipids, including cholesterol, are suppressed whereas several amino acids are activated in the fetal brain due to the absence of *Cav1*.

Mapping the deregulated lipids and amino acids to the compound database of Kyoto Encyclopedia of Genes and Genomes (KEGG) identified the associated metabolic pathways (see **Table S1**). Cholesterol metabolism, bile acid and steroid hormone biosynthesis were identified as common pathway modules associated with the deregulated lipids due to the loss of Cav1. On the other hand, alanine, aspartate and glutamate metabolism, GABA (gamma-aminobutyrate) shunt, histidine degradation, arginine and proline biosynthesis modules were commonly associated the amino acids and organic compounds that increased in the fetal brain of *Cav1*-null mice. RNA-seq analysis showed significant (false discovery rate < 0.05) differential expression of 2,218 genes associated with the impacted pathways in the fetal brain due to the loss of *Cav1* (**Table S2**). We identified metabolic pathway genes and cognate metabolites that were impacted either negatively (both downregulated) or positively (both upregulated) in the fetal brain in response to *Cav1* ablation (**Table S3**). Integrated analysis of transcriptomics and metabolomics data by applying biclustering (61) and canonical correlation methods (62) showed canonical correlation in the variation of metabolites and genes between Cav1-null and WT mice. Three clusters (CL1, CL2 and CL3) were identified (**Figure 2**). CL1 and CL2 represented variation of lipids and pathway genes that were canonically decreased in the *Cav1*-null fetal brain. CL3 represented amino acids and associated genes that were canonically increased in the *Cav1*-null fetal brain. Mutual information network analysis (63) and centrality test (64) revealed key metabolites and genes that played central roles in gene-metabolite interactions in these clusters.

**Figure 2.**
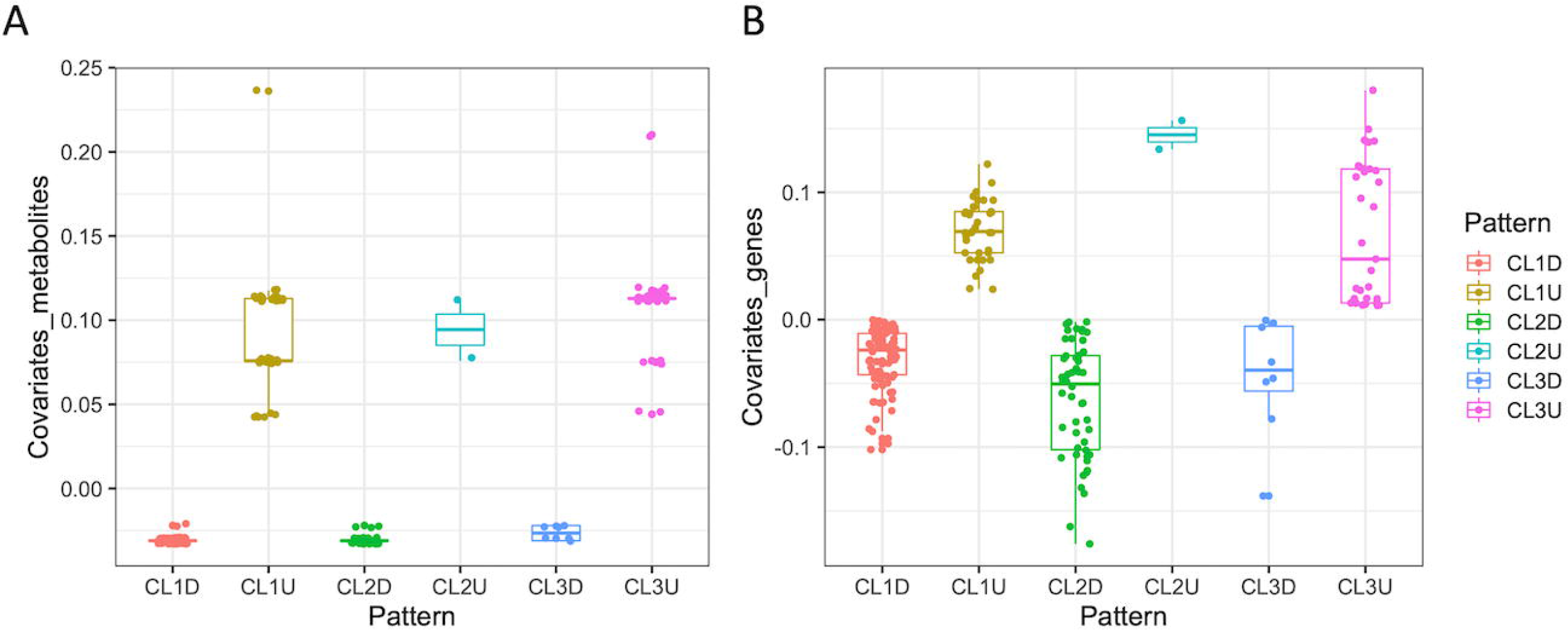
Canonical covariation of metabolites and genes in the fetal brain due to *Cav1* ablation. The box plots show covariates of metabolites (A) and associated genes (B) that were canonically either upregulated (U) or downregulated (D) with three clusters (CL1, CL2 and CL3).

Cholesterol along with *Aqp1* and *Adcy3* played central roles in the crosstalk of genes and metabolites of CL1 and CL2. Glutamic acid and *Dnah2* played similar role in the crosstalk of genes and metabolites of CL3. In addition, specific ligands and their receptors that were part of these clusters (**Table S5**), the majority of which were downregulated in the fetal brain due to *Cav1* ablation.

#### Impact of Cav1 ablation on single cells of fetal brain

Lack of membrane caveolae due to *Cav1* ablation is known to impact endothelial cell metabolism (65, 66) and vascular abnormality (67) which in turn contribute to progression of glia deregulation (68, 69), neurodegeneration and ultimately AD symptoms (70–72). Thus, we wanted to know how the different cell types of the fetal brain were impacted by *Cav1* ablation. Towards this aim, single nucleus RNA sequencing (snRNA-seq) was performed that generated expression data from 9,746 cells of WT brain and 10,753 cells *Cav1*-null fetal brain. Data analysis by *Seurat* (73) identified expression clusters of neurons, astrocytes, oligodendrocytes, ependymal, radial glia and microglial cells (**Figure 3**) based on expression of known marker genes of brain cells (74–80). Significant (*p* < 0.05) differential expression (DE) of genes was observed in individual clusters. These cluster specific markers included different metabolism genes identified from the integrated analysis metabolomics and bulk RNA-seq data. Pair-wise correlation of these metabolism marker genes among the cell types showed that cholesterol, palmitic acid and stearic acid genes were distinctly correlated among the astrocytes and microglia cells of fetal brain of Cav1-null compared to WT mice (**Figure 4**). Comparative cluster analysis further showed that *Apoe*, a well-known marker of AD (19, 81, 82), was expressed in a distinct manner in the astrocytes due to *Cav1* ablation (**Figure 5**). In the fetal brain of Cav1-null mice, *Apoe* expression showed a cluster pattern with specific lipid metabolism genes such as *Acaca, Tecr, Scd2, Fads, Sort1, Hacd3, Larpap1, Vaba* and *Lrp1*. This pattern was not observed in the WT brain suggesting that *Apoe* expression was linked to lipid deregulation of astrocytes in due the absence of *Cav1*. Furthermore, the number of glial cells expressing specific lipid metabolism genes was significantly different in *Cav1*-null compared to WT brain. The *Cav1*-null brain showed significantly more number (2×2 contingency test, p < 0.01) of astrocytes expressing *Notch2*, *Fgfr3* and *Ndufa12* than that WT brain (**Figure 6**). But oligodendrocytes expressing the same genes showed an opposite pattern. Similar difference was also observed between number of oligodendrocytes and ependymal cells expressing *Gipr*, *Aldh1b1* and *Wnt7b* in *Cav1*-null relative to WT brain (**Figure 6**).

**Figure 3.**
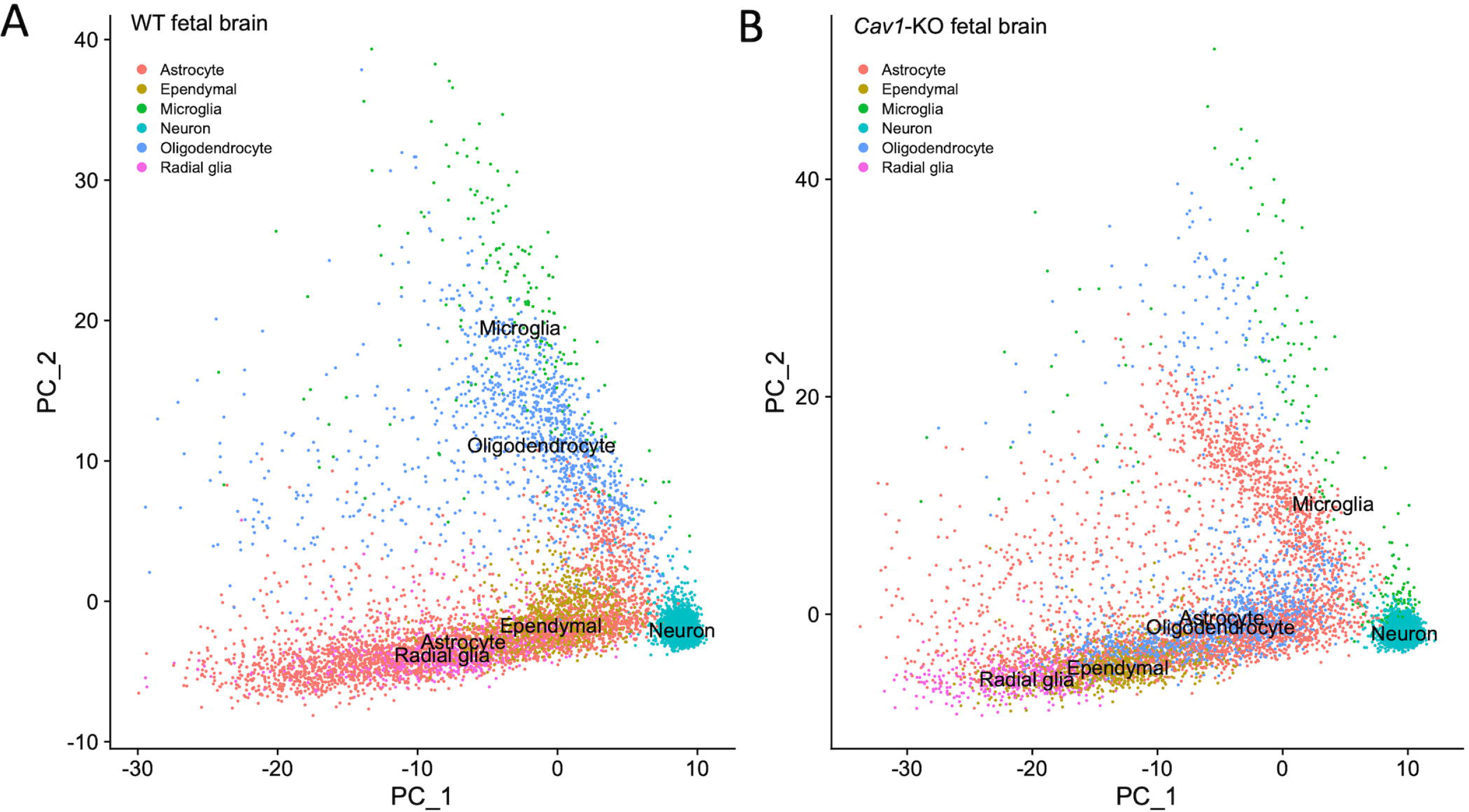
Dimensional plot of single-cell expression of WT (A) and Cav1-KO fetal brain (B). PC1 and PC2 represent the principal component axes. Cell types are color codes as shown in the legends.

**Figure 4.**
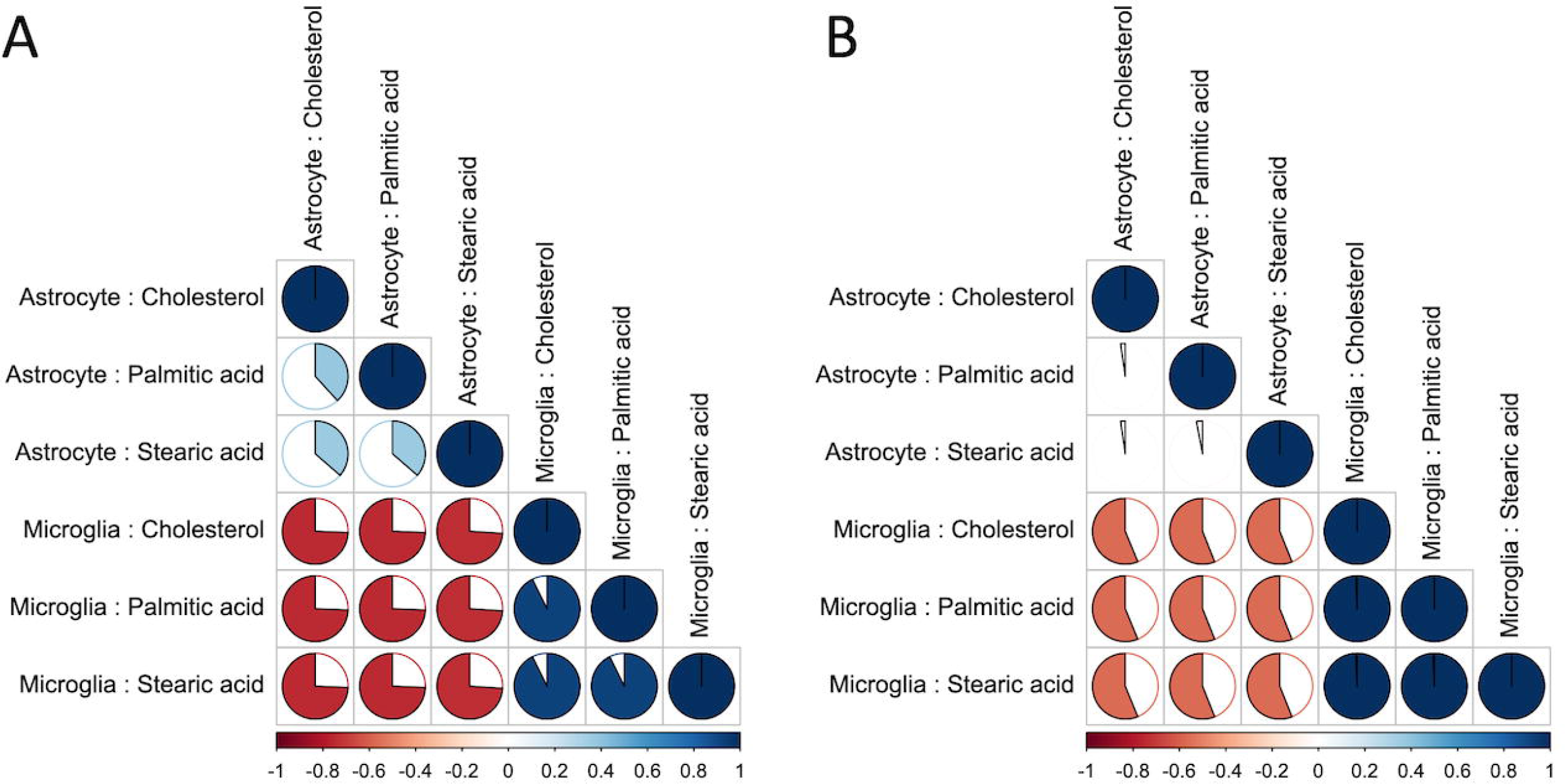
Pair-wise correlation plot showing variation in the cluster distance of gene expression associated with cholesterol, palmitic acid and stearic acid in WT (A) and *Cav1*-null (B) fetal brain. The lower panel of correlation matrix is shown in both. Color represents correlation direction and the pie represents the absolute value of correlation.

**Figure 5.**
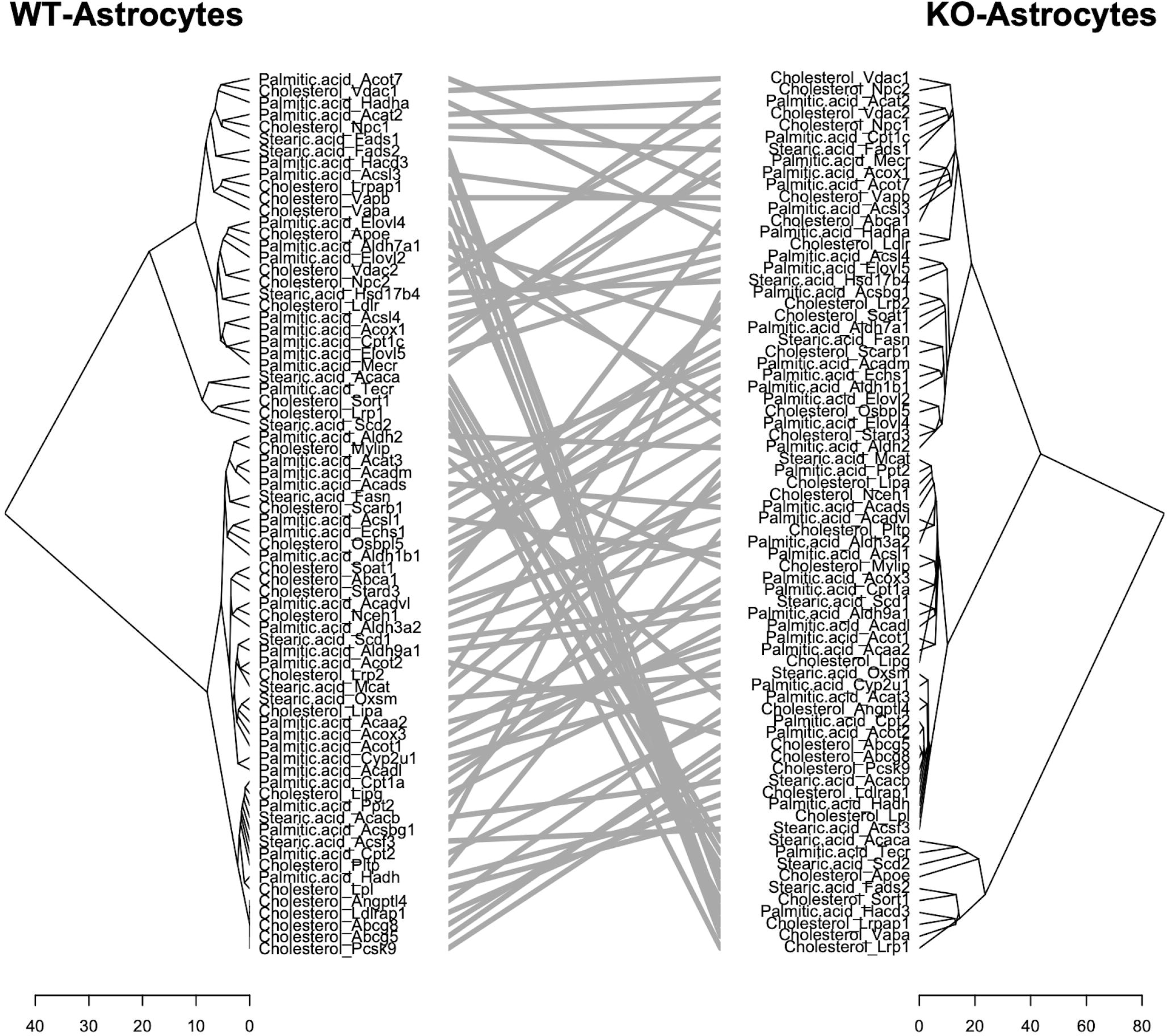
Tanglegram showing hierarchical cluster trees in a comparative manner for variation of lipids and genes associated with those lipids between WT and *Cav1*-null fetal brain. The nodes of each dendrogram represents the lipid and associated gene. Tangles show the relative positions of each node between the two dendrograms. The scale of branch lengths is shown below the dendrograms.

**Figure 6.**
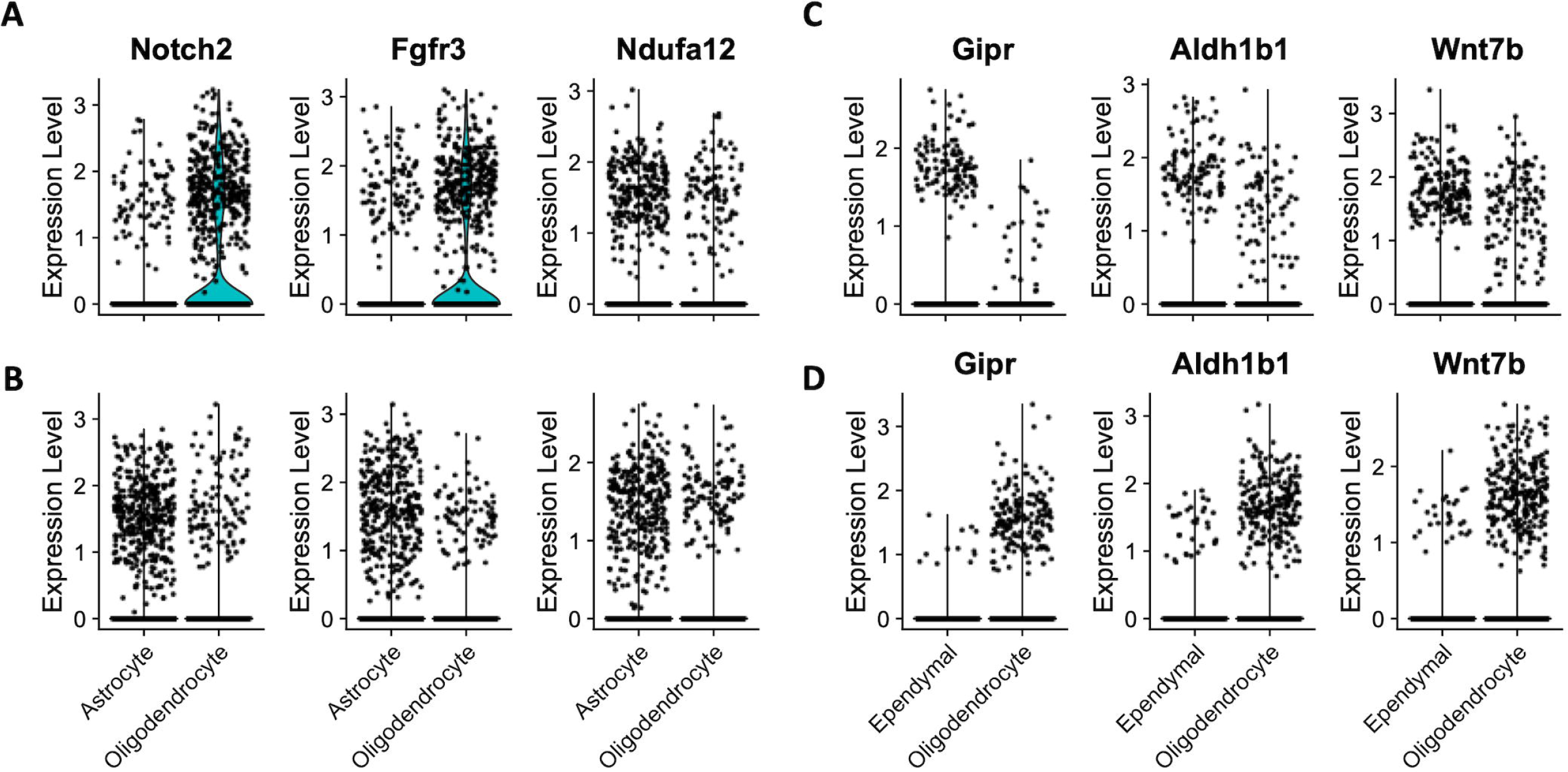
Violin plots of single cell expression of specific metabolic genes in specific types of glia cells. Each dot represents a cell. X-axis shows the cell type and Y-axis shows expression values. The kernel density, wherever applicable, is shown with color (green). **A**. Expression pattern of *Notch2, Fgfr3* and *Ndufa12* in the astrocytes of WT fetal brain. **B**. Expression pattern of *Notch2, Fgfr3* and *Ndufa12* in the astrocytes of the *Cav1*-null fetal brain. **C**. Expression pattern of *Gipr, Aldh1b1* and *Wnt7b* in the astrocytes of WT fetal brain. **D**. Expression pattern of *Gipr, Aldh1b1* and *Wnt7b* in the astrocytes of *Cav1*-null fetal brain.

#### Epigenetic modification of metabolism genes in the fetal brain due Cav1 ablation

Next, we investigated epigenetic changes of genes associated with deregulated metabolites in the fetal brain. Methylation changes of cytosine-guanine (CpG) sites associated with mouse epigenetic clock (83) were compared in the fetal brain of *Cav1*-null relative to WT mice (**Table S7**). This analysis identified CpGs (n=784) that were methylated at a higher level in the fetal brain due to Cav1 ablation (**Figure 7**). An opposite pattern was observed with a lesser number of sites (n=492). These methylations were found in specific metabolism genes which are also aging-related genes based on epigenetic clock analysis (83). The hyper-methylation sites in *Cav1*-null brain were associated with *Stard3*, *Dapk1*, *Bdnf*, and *Cpt1c* whereas the hypo-methylation sites were associated with *Wnt3a, Wnt7b, Apoe* and *Gabra5*. *Fzd2* and *Pou2f2* showed both hyper and hypo-methylation sites in the fetal brain of Cav1-null mice.

**Figure 7.**
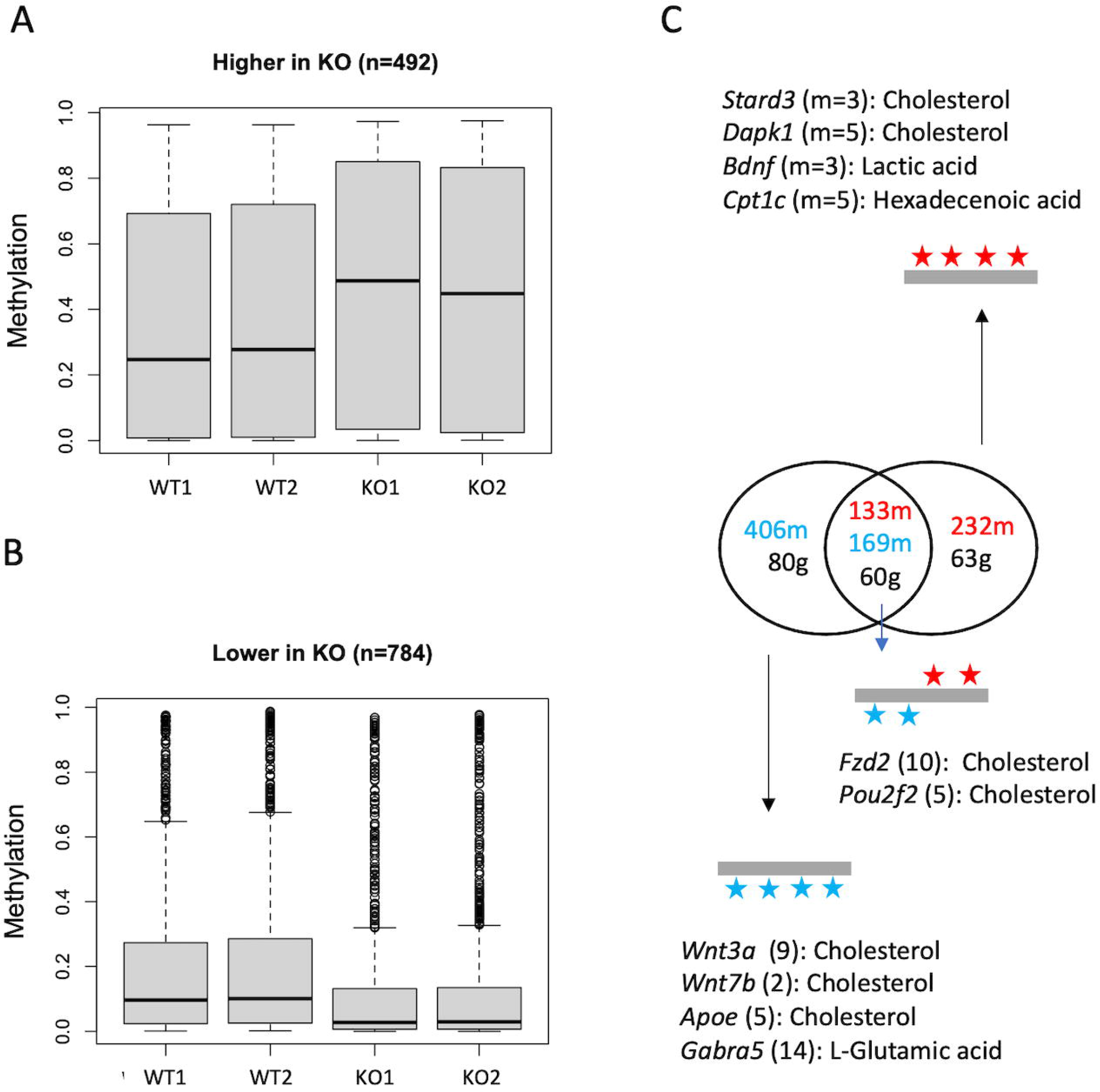
Epigenetic changes of fetal brain, and influence on gene expression of aging brain. **A**. Higher level of methylation (beta values, shown in y-axis) in *Cav1*-null brain compared to WT brain. **B**. Lower level of methylation in *Cav1*-null compared to WT brain. Brain from both sexes (female:1, male:2) are shown. **C**. Specific metabolisms genes accumulated varying number (values shown within parentheses along with metabolite name) methylation that were either increased (shown in red) or decreases (shown in blue) in the fetal brain of *Cav1*-null compared to WT mice. The Venn diagram shows that specific genes accumulated both the types of methylations.

Next, we asked how these metabolism genes epigenetically altered in the fetal brain of KO mice were impacted in the aging brain. RNA-seq was performed to determine gene expression changes of in brain of 70-weeks old *Cav1*-null compared to WT mice. Differential expression (DE) analysis identified 2,747 genes altered significantly (false discovery rate < 0.05) in the aging brain of *Cav1*-null compared to WT mice (**Table S8**). We identified genes (n=290), henceforth referred to as common DE genes (CDGs) that were significantly altered both in fetal as well as aging brain (**Table S9**). These genes were either upregulated or downregulated along with 1) genes that were associated with deregulated metabolites in the fetal brain, referred to as MDGs, and 2) genes that were epigenetically altered in the fetal brain, referred to as EDGs (**Table S10)**. A set of 345 genes were coordinately downregulated in the fetal brain due to *Cav1* ablation. These genes included 203 CDGs, 23 MDGs, and 119 EDGs. A set of 100 genes comprised of 87 CDGs, 1 EDG and 12 MDGs were coordinately upregulated in the fetal brain due to *Cav1* ablation. The same gene types, but in varying numbers, were either downregulated or upregulated in 70-weeks old brain due to the absence of *Cav1* (**Table S11**) suggesting that gene activity of aging brain of *Cav1*-null mice was associated with the metabolic and epigenetic changes in the brain at fetal stage.

## Discussion

In this study, we showed that metabolism of fetal brain was deregulated due to the absence of *Cav1*. The premise of this hypothesis is based on known function of *Cav1* in the metabolic homeostasis of cells (42). Metabolic deregulation during fetal stage can lead to neurodegenerative disorders upon aging (84–88). The developmental aging (DevAge) theory originally proposed by Dilman (89) suggests links between early development and aging (90–93). While the molecular links between early-life and adult diseases remain poorly understood, emerging evidence suggest that epigenetic memory of fetal abnormality can influence health and diseases during aging (94).

Our analysis showed that lack of *Cav1* perturbed specific lipids in the fetal brain. Lack of *Cav1* showed a reduced level of cholesterol in the fetal brain. Reduced level of cholesterol was also observed in embryonic fibroblasts and peritoneal macrophages due to lack of *Cav1* in an earlier study (42). The same study (42) also observed that reduction of cholesterol was associated with an increase of acyl-CoA:cholesterol acyl-transferase suggesting higher level of esterification of cholesterol in response to *Cav1* ablation. Caveolin-1 and cholesterol, along with sphingolipids, are integral components of caveolae. However, brain cholesterol is largely independent circulating cholesterol in the blood due blood-brain barrier. In the brain, cholesterol is primarily synthesized in the astrocytes which is supplied to the neuronal cells via lipoprotein primarily *Apoe* by the transporter *Abca1*. Within neurons, cholesterol level is controlled by activation of the oxysterol 24S-hydroxycholesterol (24S-HC) that removes excess cholesterol. In the developing mouse brain, specific nuclear receptors such as liver-X receptors (LXRs), play major roles in the regulation of neurogenesis (95). LXRs can be activated by 24S-HC in the brain (96) that results in the activation of cholesterol transporter *Abca1* and increased efflux of cholesterol from the astrocytes (97). During brain development, cholesterol plays critical roles in brain patterning, myelination, neuronal differentiation, and synaptogenesis (98) and deregulation of cholesterol in the fetal brain can lead to developmental defects of the brain (99).

Besides cholesterol, the metabolomics analysis also showed reduction of palmitic acid and stearic acid and increase of several amino acids in the fetal brain due to the absence of *Cav1*. The decrease of the fatty acids and increase of amino acids may be due to differential lipid-protein interaction of fetal brain due to the loss of *Cav1*. This supports the idea that cholesterol levels influences lipid-protein interaction whose deregulation in brain can lead to neurodegenerative diseases (100, 101). Integrative analysis of metabolomics and gene expression data further showed that specific metabolites and genes were coordinately upregulated or downregulated in the fetal brain due to the absence of *Cav1*. These genes and metabolites were commonly associated with specific KEGG pathway modules (a module represents interacting pathways) including beta-oxidation (mmu_M00086), cholesterol biosynthesis (mmu_M00101), and triacylglycerol biosynthesis (mmu_M00089). Beta-oxidation of non-esterified fatty acid is key to utilization of glucose oxidation for cellular energy instead fatty acids (98), and deregulation of triacylglycerols in the brain can lead to cognitive impairment including Alzheimer’s (102). Our data showed that ablation of *Cav1* triggered a complex gene-metabolite interactive response in the fetal brain. Several of the altered genes in the brain, due to *Cav1* ablation, were known ligand and receptors. They largely represented both multi-receptor ligands (such as apolipoprotein E, calmodulin, collagen 4a, fibroblast growth factors, jagged canonical Notch ligands) and multi-ligand receptors (such as integrins, fibroblast growth factors, Notch, adenylate cyclases) whose roles in brain diseases are well documented (103–107).

By performing snRNA-seq, we identified specific cell types in the fetal brain that were impacted due to Cav1 ablation. We applied a meta-analysis approach that identified metabolism genes differentially expressed in brain cells based on bulk RNA-seq DE genes. We have described the approach of meta-analysis of marker genes in single-cell RNA-seq data based on bulk RNA-seq data in our earlier study (108). Using this approach, we observed that the genes associated with glutamate and cholesterol were expressed in a greater number of cells of microglia, oligodendrocytes and neurons in the fetal brain of *Cav1*-null compared to WT mice. We observed that specific cell types were present in differential abundance in the fetal brain in *Cav1*-null mice. Astrocytes expressing *Notch2*, *Fgfr3* and *Ndufa12* were significantly more abundant than oligodendrocytes in the fetal brain of *Cav1*-null compared to WT mice. On the other hand, ependymal cells expressing *Gipr*, *Aldh1b1* and *Wnt7b* were significantly lower in number than oligodendrocytes in *Cav1*-null fetal brain. These findings supported the idea that *Cav1* played a role in regulating cell number by controlling cell death and proliferation due changes in either physiological or pathological conditions of the brain (109).

Finally, we investigated if genes associated with the deregulated metabolites were epigenetically altered in the brain. The rationale of this hypothesis is based on the evidences that metabolic changes alter epigenetic program of the fetal brain (32, 84, 110–115). By profiling methylation of mouse multi-tissue epigenetic clock (116, 117), our analysis showed that 50% of the CpGs associated with mouse epigenetic clock were differentially methylated in the fetal brain due to loss of *Cav1*. We wanted to know how the metabolism genes epigenetically altered in the fetal brain were expressed in the brain at old age. The WT and *Cav1*-null mice were aged to 70 weeks, and brain gene expression was profiled by RNA-seq. The data analysis showed that metabolism genes epigenetically alerted in the fetal brain influenced gene expression of the aging brain. Network analysis showed that these genes impacted gene crosstalk in the brain. This is a significant finding that supports the emerging idea that methylation changes can have long-term effects, a concept called as ‘epigenetic memory’ (26) which suggests that early-life metabolic stresses are epigenetically linked to adult health and diseases (22, 24, 26–28). In the context of epigenetic memory, our study suggests that ablation of *Cav1* elicits an epigenetic memory of the metabolic disorder of fetal brain that confounds the epigenetic clock of the brain upon aging. This is a possible mechanism of early-life links of rapid brain aging known in *Cav1*-null mice (118) which not only supports our earlier findings that brain aging originates in the fetus (21), but also suggests that epigenetic memory of fetal metabolic disorder may be linked to genesis of Alzheimer’s symptoms in early ages of these mice.

## Materials and methods

### Animal breeding and sample collection

The WT (C57BL/6J) and *Cav1*-null mice were obtained from Jackson Laboratory (stock numbers: 000664 and 007083 respectively). Adult females were mated with fertile males to induce pregnancy. The start of pregnancy (day 1) was considered when a vaginal plug was observed. The pregnant mice were euthanized on day 15 and the whole fetal brain was collected as described earlier (108). The samples were washed in sterile PBS and snap frozen in liquid nitrogen. Additionally, *Cav1*-null mice were aged to 70 weeks to collect adult brain. All animal procedures were approved by the Institutional Animal Care and Use Committee of the University of Missouri-Columbia and were conducted according to the *Guide for the Care and Use of Laboratory Animals* (National Institutes of Health, Bethesda, MD, USA).

### Metabolomics analysis of fetal brain

The WT and *Cav1*-null fetal brains in three replicates were subjected to untargeted metabolomics profiling of by gas chromatography–mass spectrometry (GC-MS). Agilent 6890 GC coupled to a 5973N MSD mass spectrometer at the University of Missouri Metabolomics Core was used in the analysis. As quality control and calculation of retention index, a standard alkane mix was used. Metabolites were identified from spectral analysis by the AMDIS (Automated Mass Spectral Deconvolution and Identification System) using an in-house database used by our Metabolomics Core. We also used a commercial NIST17 mass spectral library to search for additional metabolites that were not present in the in-house library. The abundance of the identified metabolites was determined by the Metabolomics Ion-Based Data Extraction Algorithm (MET-IDEA) (119).

### RNA-sequencing of fetal brain

Total RNA was isolated from day 15 fetal brain of WT and *Cav1*-null mice as well as from week 70 old *Cav1*-null mice using an AllPrep DNA/RNA Mini Kit (Qiagen, Cat No./ID: 80204) following the manufacturer’s instruction. Off note, RNA-seq of week 70 WT mice brain was generated from our earlier study (Accession # GSE215138). Samples were homogenized with 500 ul RLT buffer (Qiagen, Cat No./ID: 79216) supplemented with 5μl of 2-mercaptoethanol. The homogenate was transferred to a fresh tube and centrifuged for 1 minute at ≥ 8000 x g. From the supernatant, 750μl was transferred to fresh tube and mixed with 1 volume 70% ethanol to precipitate RNA. RNA was eluted in 30μl nuclease-free water twice to a total volume of 60μl. Concentration of RNA was determined using Nanodrop 1000 spectrophotometer (Thermo Fisher Scientific). RNA Integrating Number (RIN) was determined (using Agilent2100). RNA was used for preparation of libraries and RNA sequencing (RNA-seq) by Novogene Cooperation Inc (8801 Folsom Blvd #290, Sacramento, CA 95826). Each library was sequenced to 20 million paired end reads of 150 bases using a NovaSeq sequencer. RNA-seq data analysis was performed as described in our earlier works (108, 120). Briefly, the quality of raw sequences was checked with FastQC followed by trimming the adaptors from the sequence reads by *cutadapt*. The *trimmomatic* tool was used to perform base quality trimming (Phred score >30) by sliding window scan (4 nucleotides). The quality reads were then mapped to the mouse reference genome GRCm39 using Hisat2 aligner (121). Read counting from the alignment data was performed by *FeatureCounts* (122). The feature count data was then analyzed by *edgeR* (123) to determine significance of differential expression of *Cav1*-null compared to WT brain.

### Single-nuclei expression profiling of fetal brain

Single-nuclei RNA sequencing (snRNA-seq) (124) was performed to profile gene expression of single cells of day 15 fetal brain of WT and *Cav1*-null mice. Single nuclei from fetal brain were isolated using Pure Prep Nuclei Isolation kit (Sigma, St. Louis, MO, USA) as per manufacturer’s instructions. Briefly, frozen brain samples after thawing were minced into small pieces and added to the lysis buffer supplied in the kit. Using a dounce homogenizer, the sample were be homogenized till the solution looks evenly mixed, which generally required 15-20 dounces. A 70μm cell strainer was be used to filter the nuclei from the lysed cells, diluted and layered over a freshly prepared 1.8M sucrose cushion solution to collect single nuclei. After centrifugation and suspension with the provided storage buffer (ice cold), final purification of single nuclei was be performed by filtering through a 40μm cell strainer. The purified nuclei were then counted using a Countess II FL Automated Cell Counter (ThermoFisher).

The freshly prepared nuclei were used to prepare sequencing libraries using 10X Genomics Chromium Single Cell 3◻ GEM, Library & Gel Bead Kit v3.1 at the University of Missouri Genomics Technology Core. The nuclei suspension, reverse transcription master mix, and partitioning oil were loaded on a Chromium Next GEM G chip. Post-chromium controller GEMs were transferred to a PCR strip tube and reverse transcription performed on an Applied Biosystems Veriti thermal cycler at 53°C for 45 minutes. cDNA was amplified for 12 cycles and purified using Axygen AxyPrep MagPCR Clean-up beads. cDNA fragmentation, end-repair, A-tailing, and ligation of sequencing adaptors was performed according to manufacturer’s specifications. The libraries were quantified using a Qubit HS DNA kit. The fragment size was analyzed using an Agilent Fragment Analyzer system. Libraries were sequenced on an Illumina NovaSeq 6000 with a sequencing configuration of 28 base pair (bp) on read1 and 98 bp on read2.

Each library was sequenced to a depth of 20,000 paired-end (single-indexing) reads per nucleus. The base call (BCL) files were processed by Cell Ranger pipeline (v. 3.0.1) to generate the FASTQ files. The STAR aligner (125) was used to map the reads in the FASTQ files to the mouse reference genome GRCm39 to generate read count data of genes expressed in the single cells.

The count data was processed by *Seurat* (126) to identify expression clusters and assign clusters to cell types. Briefly, data integration was performed by identifying integration anchors for the first 20 dimensions of data variation among the WT and *Cav1-null* brain samples. The scaled-normalized integrated data was subjected to cluster identification by principal component analysis (PCA) and non-linear dimensional reduction by tSNE (t-distributed stochastic neighbor embedding) (127). The ‘*FindAllMarkers*’ function of *Seurat* was used to identify marker genes of each cluster. The cell types of the expression clusters were annotated based on marker genes of brain cells curated in *PanglaoDB* (128) and recently published single-cell RNA studies of brain (74–80). Comparative cluster analysis of gene expression was performed using *Dendextend*.

### DNA methylation analysis

DNA methylation was profiled for 2,045 CpG sites associated with mouse multi-tissue epigenetic clock (129) for WT and *Cav1*-null fetal brain. The epigenetic clock was analyzed as methylation changes of clock CpGs occur in correlation with age of mice (21). Methylation profiling was performed by ZYMO RESEARCH (Irvine, CA 92614, U.S.A). Briefly, DNA from frozen brain samples were purified using the Quick-DNATM Miniprep Plus kit (Cat. No. D4068). Bisulfite conversion was performed using the EZ DNA Methylation-Lightning TM Kit (Cat. No. D5030), followed by enrichment for target loci, and sequencing on an Illumina® HiSeq instrument. Data analysis was performed using *Bismark* (130) to extract methylation sites and beta-values as methylation level of each site.

### Bioinformatics and statistical analysis

The metabolites identified from the untargeted metabolomics assay of brain samples were used to identify the associated pathways by mapping to KEGG compound database. The genes associated with the identified metabolic pathways were extracted from the KEGG orthology (KO) database. The genes and metabolites mapping to the same pathways were identified which allowed to merge corresponding expression and the metabolic data. The resulting dataset was analyzed by applying two-dimensional clustering technique based on the mean squared residue method (61). In this method, variation score for each metabolite-gene pair was calculated using the equation

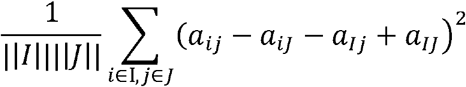

where aiJ is the mean of row i, aIj is the mean of column j, and aIJ is the overall mean of the rows and columns. The calculated scores were evaluated by fixed values of alpha (scaling parameter) and delta (maximum threshold values) using the *R* package *Biclust*. The procedure involved sequential node deletion based on the mean squared residue relative to the specified delta and alpha values (delta=.001 and alpha=1). In the first step, the rows and columns were deleted if they had scores larger than alpha times the matrix score. In the second step, the rows and columns with largest scores were removed and in the final step, rows and columns were added until the desired scaling (specified by alpha) was reached.

The combined data of genes and metabolites were subjected to mutual information (MI) network analysis (63). MIs were calculated in a pair-wise manner among the gene-metabolite pairs based on variation of RNA-seq and metabolomics data. The maximum relevance minimum redundancy (MRMR) method (131) was applied to generate weighted adjacency matrix from the MI scores. Network analysis was performed using *minet* (132). Canonical correlation analysis (CCA) was performed, as described in our earlier work (21), to determine co-variation of expression between genes and metabolites in *Cav1*-null relative to WT brain. The receptor and ligand genes were identified from database developed by Ramilowski *et al*. (133), and network analysis of expression of the ligand-receptor pairs was performed as described in our earlier work (134).

## Competing interests

The authors declare that they have no competing interests

## Funding

This research was supported in parts by funding from the start-up grant from the University of Missouri, Columbia (SKB).

## Authors’ contributions

SKB designed the study, MI performed experiments, MI and SKB performed data analysis, MI and SKB wrote the paper.

## Availability of data and material and Accession numbers

The raw and processed data of single nuclei RNA sequencing have been submitted to the Gene Expression Omnibus database under the accession number GSE214759. The raw and processed data of bulk RNA sequencing have been submitted the same database under the accession number GSE215139. The metabolomic and epigenetic data are provided as supplemental data with this manuscript.

## Acknowledgements

The authors acknowledge Nathan J. Bivens and Mingyi Zhou of the University of Missouri Genomics Core for 10x Genomics services, and Shankar P. Poudel for reading the manuscript.

## Notes

### Competing Interest Statement

The authors have declared no competing interest.

